# Genome size variability reflects contrasting ecological strategies in *Rhodoferax* and *Allorhodoferax*

**DOI:** 10.64898/2026.11.13.766366

**Authors:** Alejandro Rodríguez-Gijón, Víctor González-Mallén, Luis Miguel Rodriguez-R, Sarahi L. Garcia

## Abstract

Genome size is increasingly recognized as an important feature of aquatic prokaryotes, yet the ecological and evolutionary processes underlying this variability within freshwater bacterial lineages remain poorly understood. Here, we investigated the interplay between genome size, metabolic potential, and environmental distribution within the freshwater bacterial lineage historically classified as *Rhodoferax* (family *Burkholderaceae*) as a model system. We curated a dataset of 345 high-quality genomes grouped into 101 species-level clusters (ANI >95%) with estimated genome sizes ranging from 2.41 Mbp to 6.92 Mbp. Phylogenomic analyses revealed substantial evolutionary divergence within this lineage, supporting the distinction of *Rhodoferax* and *Allorhodoferax* and the recognition of two additional genera, which we propose to name *Quasirhodoferax* and *Pararhodoferax*. *Rhodoferax* and *Allorhodoferax* were widely distributed across global freshwater ecosystems and reached cumulative relative abundances of up to 20.4% in lake Lomtjärnan (Sweden). Species-level representatives with contrasting genome sizes exhibited distinct distributions across oxygen and depth gradients in stratified lakes, accompanied by differences in metabolic potential associated with their environmental niches. Genomic island content was also associated with genome size, although this relationship differed between *Rhodoferax* and *Allorhodoferax*, indicating that genome expansion through horizontal gene transfer may contribute differently to genome size variability across these lineages. Our study shows that genome size variability within *Rhodoferax* and *Allorhodoferax* reflects contrasting ecological strategies associated with environmental conditions, metabolic specialization, and alternative trajectories of genome evolution.

## INTRODUCTION

Genome size is linked to the ecology of prokaryotes across aquatic ecosystems (Chiriac et al., 2023; Rodríguez-Gijón et al., 2023; Swan et al., 2013). Divergent patterns of genome size in Bacteria and Archaea reflect differences in nutrient diversity and availability, and energetic constraints (Giovannoni et al., 2014; Ngugi et al., 2023; Shah et al., 2024). Moreover, differences in genome size emerge when comparing taxa across distinct aquatic environments (Rodríguez-Gijón et al., 2023) and between ecological niches within these (Salcher et al., 2019). Genome size is therefore increasingly recognized as a genomic signature of the ecological strategies through which aquatic microorganisms acquire resources, interact with neighboring organisms (Von Meijenfeldt et al., 2023), and occupy environmental niches. In this context, understanding the relationship between environmental occupation and metabolic potential of prokaryotes becomes key to understand emerging patterns of prokaryotic genome size. Consistent with this framework, the Black Queen Hypothesis states that, in conditions where the biosynthesis of key metabolites is energetically costly, genes involved in these functions might be lost leading to metabolic interdependencies between neighboring cells (Morris et al., 2012). For instance, the abundant and ubiquitous freshwater clade acI (phylum *Actinomycetota*) have highly reduced genomes below 1.5 Mbp and lack a large variety of pathways involved in the biosynthesis of essential metabolites (Garcia et al., 2013; Kim et al., 2021; Neuenschwander et al., 2018). In contrast, less abundant larger genomes may support broader metabolic repertoires and increased ecological versatility (Rodríguez-Gijón et al., 2025), enabling microorganisms to exploit more variable environmental conditions or a wider diversity of substrates. Together, these observations suggest that genome size variability may reflect different ecological strategies balancing metabolic versatility, environmental specialization, and interdependence.

Among aquatic prokaryotes, freshwater bacteria represent a particularly compelling subject of study. Although freshwater prokaryotes generally exhibit larger estimated genome sizes than brackish and marine bacteria (Rodríguez-Gijón et al., 2023), recent metagenomic surveys across global freshwater microbiomes show that highly abundant and prevalent microorganisms often possess streamlined genomes and form metabolically interdependent communities (Rodríguez-Gijón et al., 2025; Shah et al., 2024). At the same time, freshwater bacterial genera exhibit substantial differences in genome size variability with lineages containing larger average genome sizes often displaying greater variance in genome size (Rodríguez-Gijón et al., 2025). As such, genome size variability itself may represent an ecologically relevant trait linked to metabolic flexibility, habitat breadth (Bentkowski et al., 2015), and adaptive potential (Maistrenko et al., 2020). In contrast, several abundant and streamlined freshwater lineages exhibit consistently low genome size variability, potentially reflecting stronger ecological specialization and constrained functional repertoires (Kim et al., 2021; Neuenschwander et al., 2018). Together, these observations indicate that freshwater microorganisms may experience contrasting selective pressures favoring either genome streamlining and a metabolic and habitat specialization, or metabolic expansion and ecological versatility.

To explore the ecological patterns derived from genome size variability in a prokaryotic model of study, we selected the genera *Rhodoferax* and *Allorhodoferax* (family *Burkholderiaceae*, class *Gammaproteobacteria*). The genus *Rhodoferax* was described for the first time in 1991 with the isolation of *Rhodoferax fermentans* from activated sludge and ditchwater (Hiraishi et al., 1991). Since then, multiple species have been isolated from freshwater environments (Finneran, 2003; Jin et al., 2020; Kaden et al., 2014; Kato et al., 2024; Kitikhun et al., 2024; Li et al., 2020; Park et al., 2019; Zhou et al., 2019), microbial mats (Baker et al., 2017; Madigan et al., 2000), and sludge (Farh et al., 2017). This genus is characterized for a large variation in genome size, ranging from 3.67 Mbp in *Rhodoferax bucti* (Zhou et al., 2019) to 5.90 Mbp in *Rhodoferax koreense* (Farh et al., 2017). Such extensive genome size variability within a single genus provides a valuable opportunity to investigate how genome architecture relates to metabolic potential, habitat occupation, and ecological strategy while minimizing the effects of deep phylogenetic divergence. For instance, it is already known that cultured strains within the genus display broad metabolic versatility, including photosynthesis, phototrophy, chemoorganotrophy, and fermentation, as well as different levels of tolerance to oxygen (Kaden et al., 2014). These cultured species also utilize diverse carbon sources, such as cellobiose, fructose and arabinose, amongst others (Hiraishi and Imhoff, 2015; Kitikhun et al., 2024). However, a recent study exploring the cultivability of freshwater prokaryotes (Salcher et al., 2025) re-defined genus *Rhodoferax* into two diverging genera (*Rhodoferax* and *Allorhodoferax*), but the ecological and genomic differentiation between them remains underexplored.

In order to investigate the joint ecological implications of genome size variability, the environmental context and metabolic potential, we compiled a dataset of publicly available high-quality genomes from cultures and metagenomic studies classified as genera *Allorhodoferax* and *Rhodoferax* (class *Gammaproteobacteria*). More specifically, in this study we aim: i) to review the genomic and ecological diversity within these genera, ii) to study how the interplay between metabolic potential, genome size variability and environmental context influences the relative abundance of different lineages of freshwater *Allorhodoferax* and *Rhodoferax*, and iii) to investigate the main drivers of genome size differences within the genera.

## RESULTS AND DISCUSSION

### The genera *Rhodoferax* and *Allorhodoferax* encompass substantial genetic and ecological diversity

Our study compiled 345 high-quality genomes (completeness >90% and contamination <5%; **Figure S1A**) of the freshwater bacterial lineage historically classified as *Rhodoferax.* We dereplicated all high-quality genomes into 101 species-level clusters according to an ANI (Average Nucleotide Identity) similarity threshold of >95% (Jain et al., 2018), with one representative genome of highest quality each (**Table S1**).

According to the GTDB classification r232, three species-level clusters were assigned to the genus-level clades ‘Rhodoferax_A’ and two as ‘Rhodoferax*_*B’ (**Table S1**). This taxonomic separation from genus *Rhodoferax* suggests diverging evolutionary histories. While phylogenies based on 16S rRNA gene place the type strains of ‘Rhodoferax_A sediminis’ and ‘Rhodoferax_B koreensis’ within the *Rhodoferax* genus (Farh et al., 2017; Jin et al., 2020), more recent phylogenomic analyses using 120 conserved marker proteins indicate that these two species could potentially form a monophyletic sister group to the core *Rhodoferax* lineage (Kato et al., 2024). To resolve this taxonomic ambiguity, we inferred a maximum-likelihood (ML) phylogeny using the 101 species-clusters representative genomes alongside 804 reference genomes from other genera within the same taxonomic class (**Table S2**). Our results show that ‘Rhodoferax_A’ and ‘Rhodoferax_B’ do not form a monophyletic cluster with the genomes classified as genus *Rhodoferax*, but instead constitute independent clusters (**Figures 1A** and **S2**). This taxonomic divergence is further supported by the Average Amino acid Identity (AAI; **Figure 1B**). Integrating the results provided by the ANI and AAI analyses, the ML phylogeny and the GTDB-tk classification, we conclude that a formal taxonomical revision of the genus-level clades Rhodoferax_A and Rhodoferax_B is necessary to accurately reflect the evolutionary history of these lineages. We here propose the re-naming of genus-level cluster ‘Rhodoferax_A’ to genus *Quasirhodoferax* (type species: *Quasirhodoferax sediminis*) and genus-level cluster ‘Rhodoferax_B’ to genus *Pararhodoferax* (type species *Pararhodoferax koreensis*). This new taxonomical naming will be used hereafter, and their taxonomic description can be found in the **Supplementary Discussion 1**.

**Figure 1.**
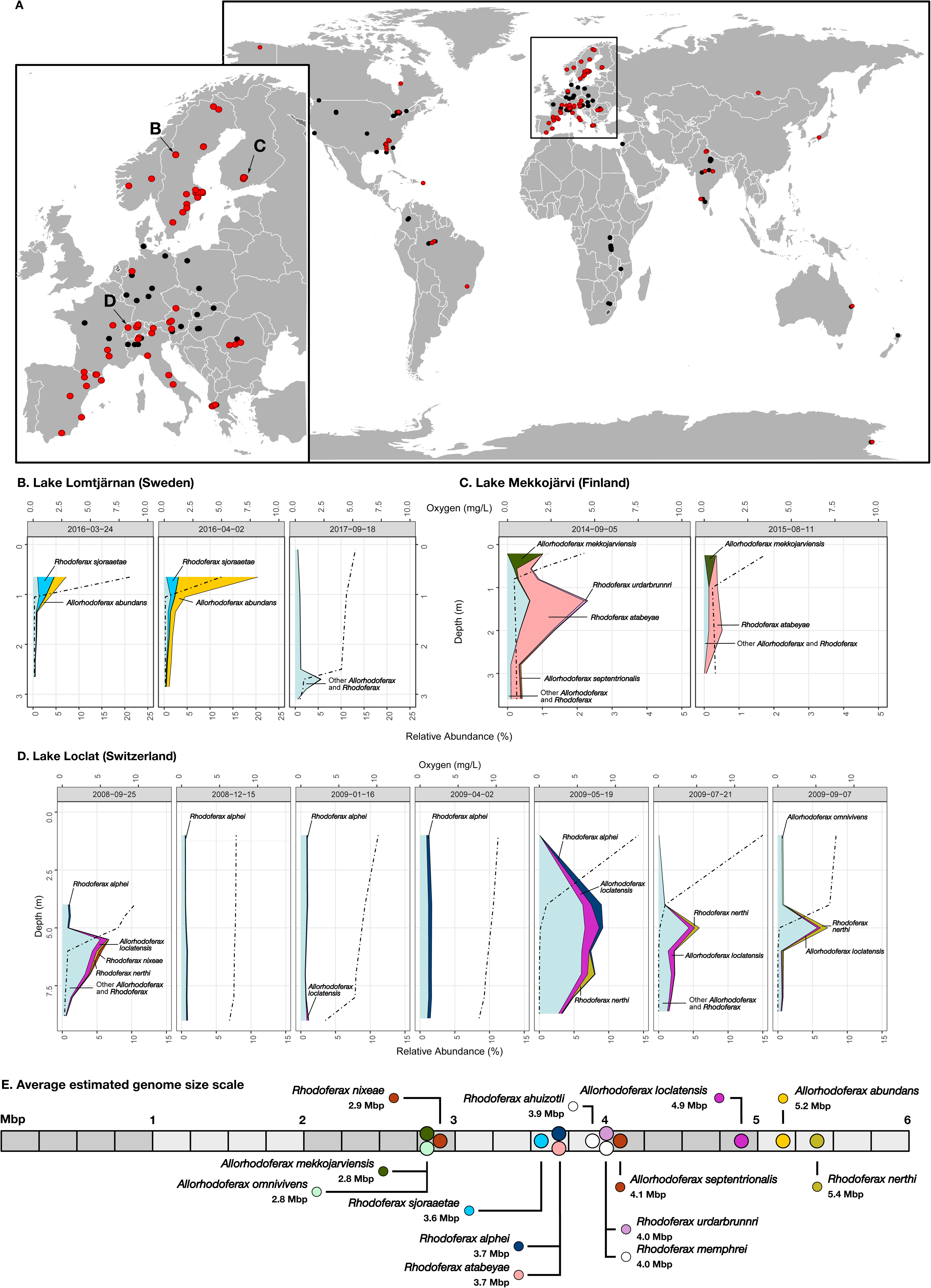
Evolutionary overview of genera *Allorhodoferax* and *Rhodoferax*. **A)** Maximum-likelihood (ML) phylogeny, following a LG+F+I+I+R10 model of evolution, of all 345 high-quality genomes (completeness >90% and contamination <5%) classified as genus *Allorhodoferax* and *Rhodoferax*, and the newly-defined genera *Quasirhodoferax* and *Pararhodoferax*. Circles on branches indicate a bootstrap support ≥90%. Information about the number of genomes per species, geographic location and type of environment of origin, and type of genome (isolate or MAG) is indicated at the end of each leaf, following the legend on the bottom-right of the figure. Expanded ML tree, including root and all outgroup genomes from class *Gammaproteobacteria*, can be found in **Figure S2**. **B)** Average Amino acid Identity (AAI) clustering, highlighting the coherent groups with the ML phylogeny. **C)** Statistical overview of the estimated genome size, GC composition and coding density across AAI clusters.

Of the remaining 96 species-level clusters, 41 are part of genus *Rhodoferax* and 55 are part of genus *Allorhodoferax*. This taxonomical differentiation is reflected statistically in genomic differences between genera, since genus *Rhodoferax* holds smaller genomes (mean = 4.10 Mbp) with lower coding densities (mean = 91.60%) and guanine-cytosine content (GC, mean = 59.34%) than genus *Allorhodoferax* (mean estimated genome size = 4.23 Mbp, mean coding density = 92.26%, mean GC = 60.90%) (**Figures S1B-S1D**). Yet, species-level clusters within both genera are notably larger than most freshwater *Pseudomonadota* genomes, averaging 3.61 Mbp (Rodríguez-Gijón et al., 2025). All 96 species-level clusters are grouped into eight differentiated clusters robustly supported by AAI and ML (**Figure 1A** and **1B**). Yet, a ninth cluster supported both by AAI and ML emerges in the picture (indicated as “cluster 8” in **Figure 1A** and **1B**). All five genomes within this cluster are classified as genus CAIKVZ01, and were originally included as reference genomes in our phylogenomic studies (**Table S2**). These genomes were assembled and binned from metagenomic samples collected in freshwater lakes in northern Europe (Buck et al., 2021). The consistent ecological and geographical distribution of the CAIKVZ01 genomes relative to the genera *Rhodoferax* and *Allorhodoferax* potentially suggests a recent evolutionary differentiation between these three genera.

Five of the AAI-clusters (i.e., 3, 4, 6, 9 and 10) consist exclusively of genomes retrieved from freshwater environments (**Figure 1A**). Despite the recent taxonomical differentiation between genera *Rhodoferax* and *Allorhodoferax* (Salcher et al., 2025), this ecology is consistent with previous amplicon-based and isolate-based surveys, which characterized *Rhodoferax* as a predominantly aquatic genus (Čuperová et al., 2013; Park et al., 2019; Petersen et al., 2025; Salka et al., 2011; Waidner and Kirchman, 2008; Zhou et al., 2019). Yet, three AAI-clusters (5, 7 and 11) reveal a broader ecological range: members of these broader ecological clusters were found in terrestrial ecosystems (such as permafrost soils, glacier sediments and forest soils), and host-associated environments (*Asterionella formosa* diatoms) (**Figure 1A** and **Table S1**). Notably, despite the highly streamlined genomes typically observed in aquatic prokaryotes (Giovannoni et al., 2014; Rodríguez-Gijón et al., 2022), freshwater *Rhodoferax* and *Allorhodoferax* clusters do not differ significantly from more ecologically diverse clusters in terms of genome size, coding density or GC composition (**Figure 1C**). These results indicate that the expansion of *Rhodoferax* and *Allorhodoferax* beyond freshwater environments is not accompanied by broad shifts in genome size, coding density or GC composition, suggesting that ecological differentiation within these genera cannot be explained by these genome architectural traits alone.

In conclusion, all our results together indicate that *Rhodoferax* and *Allorhodoferax* comprise two genetically and ecologically diverse groups of freshwater bacteria spanning distinct environmental distributions while maintaining relatively similar genome architectural characteristics. This combination of ecological diversification and broad genome size variability positions *Rhodoferax* and *Allorhodoferax* as powerful models to investigate how metabolic potential, habitat occupation, and genome architecture interact across freshwater ecosystems.

### Genome size variability reflects the interplay between metabolic potential and biotope

To explore genome size variability within genera *Allorhodoferax* and *Rhodoferax*, we first focused on its relationship with the metabolic potential of the genera. First, all high-quality genomes were metabolically annotated to predict the completeness of biosynthetic pathways for both amino acids and vitamins. Our analyses revealed a weak but statistically significant positive correlation between the estimated genome size and the average completeness of metabolic pathways involved in the biosynthesis of amino acids (*Allorhodoferax*: R = 0.14, p = 0.047; *Rhodoferax*: R = 0.18, p = 0.035) and vitamins (*Allorhodoferax*: R = 0.27, p = 5.9e-05; *Rhodoferax*: R = 0.22, p = 0.01) (**Figure S3**). Despite the limitations that can be found in the prediction of auxotrophies from genomes of non-cultivated microorganisms (Pacheco-Valenciana et al., 2025), our results align with previous studies where a positive correlation between the average module completeness and prokaryotic genome size is observed (Hessler et al., 2023; Jaffe et al., 2020; Ramoneda et al., 2023; Rodríguez-Gijón et al., 2025). However, the relatively weak correlation indicates that biosynthetic potential for essential metabolites explains only a limited fraction of the observed genome size variability, suggesting that additional metabolic and ecological factors contribute to differences in genome size.

To integrate environmental information with metabolic potential, we subset the relative abundance and prevalence estimations of all species classified as *Rhodoferax*, *Allorhodoferax*, *Quasirhodoferax* and *Pararhodoferax* from previous work (Rodríguez-Gijón et al., 2025) (**Tables S3** and **S4**). Our results show that *Quasirhodoferax* was not detected in any of the 636 freshwater metagenomes, while *Pararhodoferax* only in 3 of them located in Sweden and Greece (**Supplementary Discussion 1**). On the contrary, *Rhodoferax* and *Allorhodoferax* were jointly detected in 65.1% of the freshwater metagenomes (i.e., 414 out of 636), spanning sampled freshwater systems across multiple continents (**Figure 2A**).

**Figure 2.**
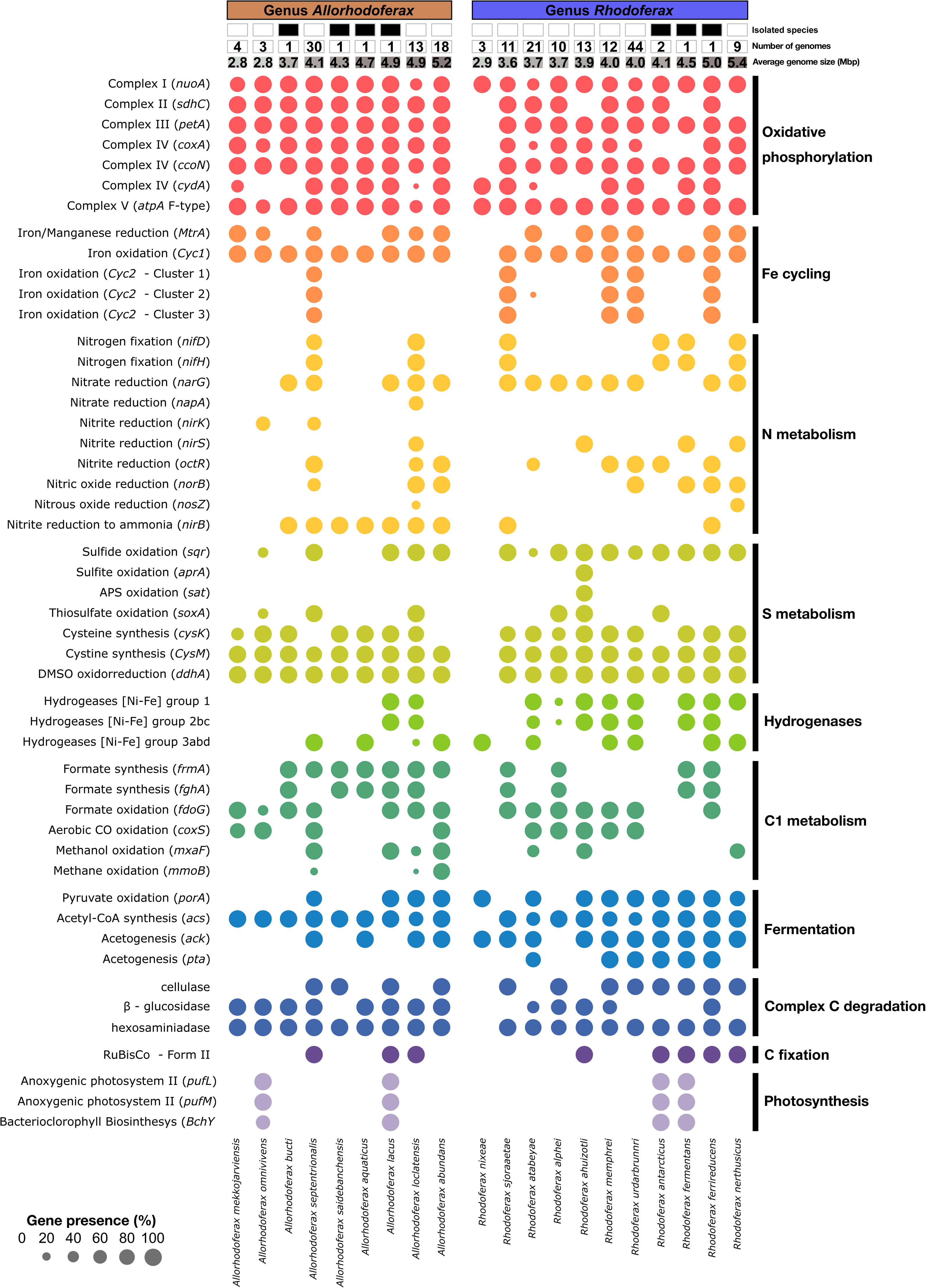
Distribution and relative abundance of genera *Allorhodoferax* and *Rhodoferax* across global freshwater bodies. **A)** Presence (red dots) and absence (black dots) of these genera across global freshwater bodies, with an emphasis on European lakes (subplot). **B-D)** Relative abundance of *Allorhodoferax* and *Rhodoferax* species-clusters across depth profiles of lakes Lomtjärnan (Sweden, **B**), Mekkojärvi (Finland, **C**) and Loclat (Switzerland, **D**). **E)** Overview of the average estimated genome size of the 13 environmental species-clusters of interest. White-colored dots refer to *Rhodoferax ahuizotli* and *Rhodoferax memphrei*, indicating their absence in the studied lakes.

However, because the metagenomic dataset is geographically biased toward Europe and North America, the absence of detections in Africa should not be interpreted as evidence that these genera are absent from African freshwater ecosystems. Their broad distribution and high prevalence nevertheless support a cosmopolitan freshwater ecology.

More specifically, the most prevalent species in our dataset (i.e., *Rhodoferax* 0967) was present in 24.06% of the metagenomes (i.e., 153 out of 636). On average, 8.22 species of *Rhodoferax* and *Allorhodoferax* are detected per metagenome when excluding the metagenomes where no species were detected, and only 16 out of the 96 species were not detected in any metagenome, including the laboratory-grown *Rhodoferax ferrireducens* (**Table S4**). The average cumulative relative abundance of these two genera was 0.96% across the 414 metagenomes where they were detected (Rodríguez-Gijón et al., 2025).

Because of the large number of *Rhodoferax* and *Allorhodoferax* species-clusters detected in our genomic dataset (n = 80), we decided to focus on the seven isolated species as well as the ten species-clusters with the most MAGs (i.e., *Rhodoferax* 0008, 0018, 0068, 0112, 0130, 0156, and 0211, and *Allorhodoferax* 0016, 0043 and 0339; **Table S1**). As these 17 species-clusters have a relatively large estimated genome size (over 3.6 Mbp), we also included in our analysis all species-clusters with a mean estimated genome size below 3Mbp that included at least 3 MAGs (i.e., *Rhodoferax* 0060, and *Allorhodoferax* 0505 and 0744) (**Table S1**). We explored the relative abundance of these 20 species-level clusters across depth and oxygen profiles in the three lakes from the Stratfresh database (Buck et al., 2021) where they showed the largest relative abundance ranging from 2.7% to 20.4% (**Figures 2B-2D**). Furthermore, we propose a new formal taxonomical naming to the 13 species-clusters recovered from MAGs (**Supplementary Discussion 1**), and this naming will be used hereafter.

In lake Lomtjärnan (Sweden), the species *Allorhodoferax abundans* and *Rhodoferax sjoraaetae*, with an average estimated genome size of 5.2 Mbp and 3.6 Mbp respectively, were the most abundant species in the lakés epilimnion, where oxygen concentration is higher (**Figure 2B** and **2E**). In the case of *Allorhodoferax abundans*, it represents up to ∼17.3% of the total microbial community above the oxycline (**Figure 2B**), potentially reflecting a seasonal bloom. This preference for oxygenated zones of the lake is reflected by their metabolic potential, since both species encode all genes involved in aerobic respiration (**Figure 3**). Yet, the presence of different genes involved in anaerobic processes such as fermentation (*ack* and *acs*) and denitrification indicate that, despite the preference of *Allorhodoferax abundans* and *Rhodoferax sjoraaetae* for oxygenated waters, they could be facultative anaerobes.

**Figure 3.**
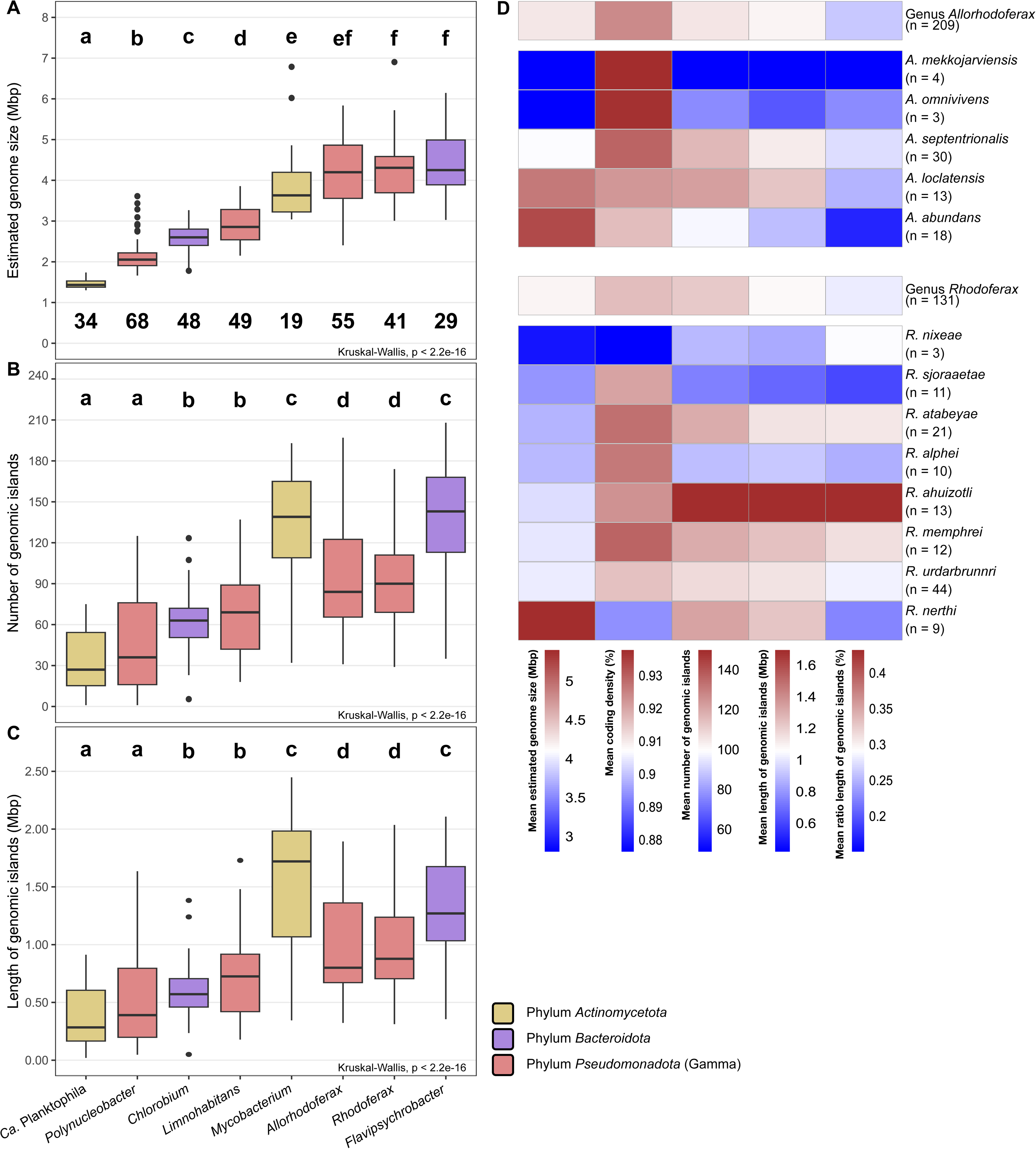
Metabolic potential of 13 selected species-clusters and 7 isolated species of genera *Allorhodoferax* and *Rhodoferax*. Boxes on the top indicate the number of genomes per species-clusters, as well as the average estimated genome size (Mbp). Circles indicate presence of a gene, size represents the percentage of the genomes that have that specific gene (following the gradient scale on the bottom-right part of the figure), and colors indicate the different metabolic categories (indicated on the right side of the figure).

In lake Mekkojärvi (Finland), the species *Allorhodoferax mekkojarviensis* was the most abundant species above the oxycline (**Figure 2C**), and holds one of the smallest average estimated genome sizes in our dataset (2.8 Mbp; **Figure 2E**). Moreover, its absence in non-oxygenated zones of the lake (below 1 meter of depth) and the lack of genes involved in anaerobic reactions (**Figure 3**), indicate a high speciation for oxygenated epilimnion waters. On the contrary, *Allorhodoferax septentrionalis* (4.1 Mbp) and *Rhodoferax urdarbrunnri* (4.0 Mbp) and *Rhodoferax atabeyae* (3.7 Mbp) have larger genome sizes (**Figure 2E**), are more abundant below the oxycline (**Figure 2C**), and have a larger repertoire of genes than *Allorhodoferax mekkojarviensis*, including genes such as the *cyc2* (oxidation of Fe^2+^), *soxA* and *sqr* (sulfur oxidation), and *ack* (acetogenesis), among others (**Figure 3**). The presence of these three species not only coincide with oxygenation, but also with Fe^2+^ concentrations: surface samples (below 1 m depth) have a concentration of Fe^2+^ ranging from 11.6 mg/L to 125.9 mg/L, while samples taken over 2 m depth have Fe^2+^ concentrations of up to 389.3 mg/L (Buck et al., 2021). Together, these patterns suggest that genome size differences among co-occurring *Rhodoferax* and *Allorhodoferax* species are associated with contrasting metabolic strategies along the oxygen and Fe² gradients of Lake Mekkojärvi.

In lake Loclat (Switzerland), none of the 20 species-clusters selected represent the most abundant species (**Table S4**). We observe that *Rhodoferax alphei* and *Allorhodoferax omnivivens* (average estimated genome size of 3.7 and 2.8 Mbp, respectively) appeared to be more abundant above the oxycline, while *Rhodoferax nixeae* (2.9 Mbp) and *Rhodoferax nerthi* (5.4 Mbp) and *Allorhodoferax loclatensis* (4.9 Mbp) were more successful below the oxycline (**Figure 2D**). This distribution is consistent with differences in the predicted metabolic potential of the five species. The species with a larger average estimated genome size (i.e., *R. nixeae* and *A. loclatensis*) are found below the oxycline (**Figure 2D** and **2E**) and encode for genes involved in the complete denitrification pathway and nitrogen fixation, such as the gene clusters *narGH* for nitrate reduction, the *nirS* for nitrite reduction, the *norB* for nitric oxide reduction, and the *nosZ* for nitrous oxide reduction (**Figure 3**). These metabolic pathways are typically associated with oxygen-limited or anoxic conditions (Jetten, 2008), and are almost absent in the species-clusters from upper layers of the lake and with more reduced genome sizes (i.e., *Rhodoferax* 0112 and *Allorhodoferax* 0744; **Figure 2D, 2E** and **3**). A similar pattern was observed in the Baltic Sea, where bacterial MAGs with bigger estimated genome sizes have a larger number of module steps involved in nitrogen metabolism (Rodríguez-Gijón et al., 2023). Similarly, these patterns are also observed at lower scale, such as the marine *Trichodesmium* spp. (phylum *Cyanobacteriota*), where nitrogen-fixing lineages contain larger genomes than non-fixing lineages (Bergman et al., 2013).

Altogether, our results suggest that genome size variability within *Rhodoferax* and *Allorhodoferax* reflects alternative ecological strategies shaped by oxygen availability, nutrient complexity and availability, and metabolic specialization across freshwater ecosystems. Rather than representing random genomic divergence, differences in genome architecture are consistently associated with niche occupation and environmental adaptation within these genera.

### Genomic islands (GIs) are associated with genome size variability

Genomic islands (GIs) are DNA segments frequently acquired through horizontal gene transfer (HGT) and are of eco-evolutionary importance given their tight link with genome size variation, potentially conferring the receiving bacterium a range of ecological and metabolic benefits (Coleman et al., 2006; Dobrindt et al., 2004). Given their association with larger genome sizes, expanded metabolic repertoires, and ecological occupation, we next explored whether the acquisition of GIs contribute to genome size variability within the genera *Allorhodoferax* and *Rhodoferax*. Hence, we explored the importance of GIs in the interplay between prokaryotic genome size, metabolic potential and ecology in our dataset of high-quality genomes, and compared with six other bacterial genera present in freshwater bodies. These genera are *Mycobacterium* and *Ca.* Planktophila (phylum *Actinomycetota*), *Chlorobium* and *Flavipsychrobacter* (phylum *Bacteroidota*), and *Limnohabitans* and *Polynucleobacter* (phylum *Pseudomonadota*) (**Table S5**) (Garcia et al., 2021; Newton et al., 2011; Nuy et al., 2020; Rodríguez-Gijón et al., 2025).

We found that the genera *Flavipsychrobacter*, *Allorhodoferax*, *Rhodoferax* and *Mycobacterium* not only have the largest estimated genome sizes in our dataset (mean >3.90 Mbp; **Figure 4A**), but they also contain the highest number and total length of predicted GIs (**Figures 4B** and **4C**). This pattern is consistent with previous observations stating that HGTand GIs are more prevalent in organisms with larger genomes (Cordero and Hogeweg, 2009), including those derived from prophages (Touchon et al., 2016). For instance, a recent genomic survey highlighted that GIs involved in pathogenicity and antibiotic resistance in *Mycobacterium avium* are essential for its environmental adaptation (Banerjee and Sur, 2025). Furthermore, the statistically positive correlation between the estimated genome size and both the number and total length of GIs observed in *Polynucleobacter*, *Chlorobium*, *Allorhodoferax* and *Flavypsychrobacter* (**Figure S5**) supports this hypothesis. Yet, we did not observe a statistical correlation between both variables in *Mycobacterium* (**Figures S5E** and **S5M**), potentially result of the relatively low number of genomes included in our study, similarly to *Rhodoferax* (**Figures S4G** and **S4O**). In the latter case, the absence of a positive statistical correlation between the estimated genome size and both the number and total length of GIs could be the result of species-specific deviations from this trend. For instance, *Rhodoferax atabeyae* and *Rhodoferax ahuizotli* hold comparatively small genomes but encode a large number of GIs spanning over large lengths (**Figure 4D**). Specifically, GIs represent up to 40% of the total estimated genome size of *Rhodoferax ahuizotli*, a large proportion when compared with marine bacteria, where this ratio is as high as 12% (Fernández-Gómez et al., 2012).

**Figure 4.**
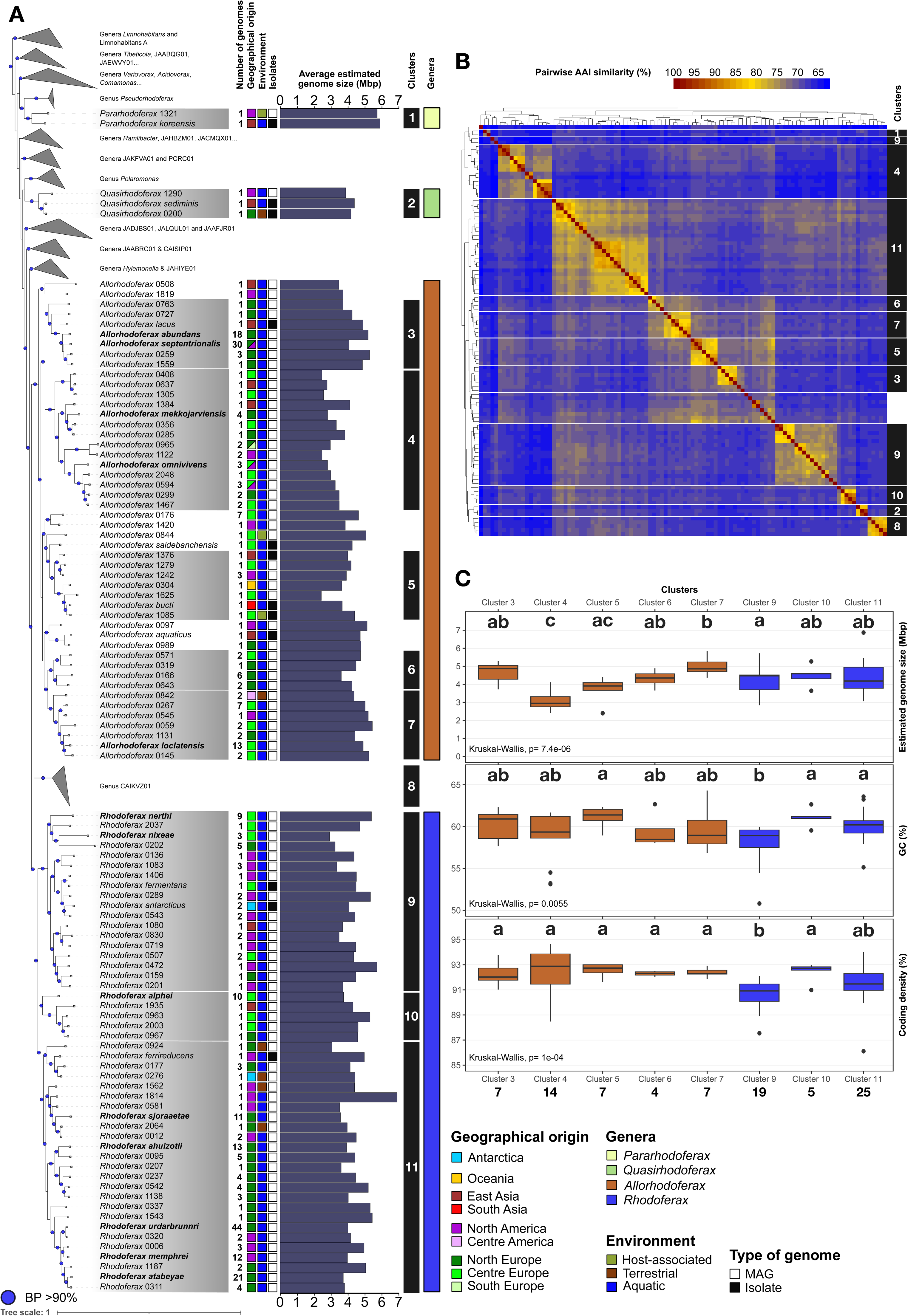
Contribution of genomic islands (GIs) to genome size variability. **A-C)** Statistical overview of the estimated genome size, number and total length of GIs across eight freshwater bacterial genera, including *Allorhodoferax* and *Rhodoferax*. Different letters indicate statistical differences (p<0.05), color of boxplots indicate phyla following the legend on the bottom of the figure, and numbers indicate the number of species-cluster representative genomes included per genus. **D)** Overview of the mean estimated genome size (Mbp), coding density (%), number of GIs, total length of GIs (Mbp) and ratio of GIs (%) across *Allorhodoferax* and *Rhodoferax* and its species-level clusters of interest.

Overall, our results reveal that predicted GI content is associated with genome size variability across several freshwater bacterial genera, including *Allorhodoferax*, but that this relationship is not universal. Within *Rhodoferax*, species-specific differences in GI content can account for substantial fractions of individual genomes without scaling consistently with total genome size. Determining the functional composition of these regions will be necessary to establish whether GI acquisition contributes directly to the metabolic and ecological differentiation observed among *Rhodoferax* and *Allorhodoferax* species.

## CONCLUSION

Our study demonstrates that genome size variability within closely related freshwater bacteria reflects distinct ecological and metabolic strategies. Across genera *Rhodoferax* and *Allorhodoferax*, differences in genome size were associated with variation in metabolic potential and with the distribution of species across environmental gradients. At the same time, the variable contribution of genomic islands among lineages highlights that genome diversification may arise through different evolutionary processes.

These findings also raise a broader question about the evolutionary significance of genome size variability itself. While highly streamlined freshwater lineages occupy a relatively narrow range of genome sizes, potentially reflecting evolutionary strategies that have already optimized genome architecture toward efficient growth under particular ecological conditions, lineages such as *Rhodoferax* and *Allorhodoferax*, which encompass several megabases of genome size variation, reveal a much broader range of genomic and metabolic variability. The evolutionary origin that resolves why some bacterial lineages retain such extensive genomic flexibility while others converge toward highly constrained genome architectures remains unresolved. Understanding the evolutionary and ecological processes that generate, maintain, or constrain genome size variability may therefore be as important as understanding genome size itself.

## MATERIAL AND METHODS

### Collection of Rhodoferax and Allorhodoferax genomes

We compiled a total of 792 publicly available *Rhodoferax* genomes from 46 different BioProjects in NCBI, including metagenome-assembled genomes (MAGs) and genomes from isolates, using the Datasets CLI tools v14.7.0 (https://github.com/ncbi/datasets). Because all these genomes are a subset from a previous study (Rodríguez-Gijón et al., 2025), the more recently described taxa within genus *Rhodoferax* have not been included in this study (Kato et al., 2024; Kitikhun et al., 2024; Petersen et al., 2025). All genomes were taxonomically re-classified using GTDB-tk v2.1.1 (Chaumeil et al., 2020) according to the GTDB classification (data version r232) (Parks et al., 2020). The quality of the genomes was assessed using CheckM v1.2.2 (Parks et al., 2015), following the typical workflow (*lineage_wf)* with default parameters. All 792 genome are of medium-to-high quality (completeness >50% and contamination <5%), of which 345 genomes were of high-quality (completeness >90% and contamination <5%) (**Figure S1**). Genome statistics and GTDB taxonomy can be found in **Table S1**.

To estimate the genome size, we divided the bin size by the completeness provided by CheckM ranging from 0 to 1. Since the correlation between the estimated genome size and the number of scaffolds for the 345 high-quality genomes was not statistically significant (**Figure S5**), we conclude that genome fragmentation does not bias our view on the variability of estimated genome size, and if so, this bias would be minimal. All 345 high-quality genomes (completeness >90% and contamination <5%) were then dereplicated using mOTUlizer v0.3.2, using an ANI threshold superior to 95% as estimator of species-level clusters (Buck et al., 2022; Jain et al., 2018). In total, we detected 101 species-level clusters from which three were classified as genus-level clade ‘Rhodoferax_A’ (re-defined as genus *Quasirhodoferax*) and two species-clusters were classified as genus-level clade ‘Rhodoferax_B’ (re-defined as genus *Pararhodoferax*), and we selected the genome with the highest quality as the representative genome of each species-cluster (**Table S1**).

### Phylogenomic analyses

Species evolutionary relationships were predicted for the 101 species classified as *Rhodoferax*, *Allorhodoferax*, *Quasirhodoferax* and *Pararhodoferax* (**Table S1**), and including 801 species representative genomes covering all genera within the family *Burkholderiaceae* and family-level designations “Burkholderiaceae_A-E” (GTDB classification r232; **Table S2**) (Parks et al., 2020). We also selected the representative genomes of the species *Halomonas olivaria*, *Marinobacter salarius* and *Pseudomonas aeruginosa* (order *Pseudomonadales*) to root the tree. We inferred a phylogenetic tree using GToTree v1.7.00 (Lee, 2019), leveraging a set of 172 gammaproteobacterial single-copy marker genes (SCGs). This pipeline predicted genes using Prodigal v2.6.3 (Hyatt et al., 2010), identified the SCGs using HMMER3 v3.3.2 (Eddy, 2011), and aligned them individually using Muscle v5.1.linux64 (Edgar, 2022). The final alignment was trimmed using TrimAl v1.4.1 (Capella-Gutiérrez et al., 2009). Later, all 172 SCGs were concatenated into an alignment matrix and inferred a maximum-likelihood (ML) phylogeny using a LG+F+I+I+R10 model of evolution and 1000 ultrafast bootstraps using IQ-TREE v2.2.0.3 (Minh et al., 2020). The best-fit model of evolution was chosen according to the Bayesian Information Criterion (BIC). Lastly, the tree was visualized using Interactive Tree of Life (iTOL) v6 (Letunic and Bork, 2024).

### Prediction of genomic islands (GIs)

The number and total length of the GIs of all 340 high-quality *Rhodoferax* and *Allorhodoferax* genomes (excluding those classified as ‘Quasirhodoferax’ and ‘Pararhodoferax’) were predicted using Alien Hunter v1.7 (Vernikos and Parkhill, 2006). This software leverages variable order motif distributions to detect variability in codons, amino acid bias and dinucleotide composition, and classifies those distinct regions from the rest of the genome as GIs. We then extracted the number and total length of the GIs, and calculated the ratio of the genome represented by these GIs (%) by dividing the total length of the GIs by the estimated genome size of the host genome.

To evaluate the importance of GIs in these genera, we also leveraged the species-representative genomes of six other bacterial genera present in aquatic environments (Rodríguez-Gijón et al., 2025) that met the same high-quality thresholds: genera *Limnohabitans* and *Polynucleobacter* (class *Gammaproteobacteria*), *Chlorobium* and *Flavipsychrobacter* (phylum *Bacteroidota*), and *Ca.* Planktophila and *Mycobacterium* (phylum *Actinomycetota*). Genome statistics were predicted using CheckM v1.2.2 (Parks et al., 2015), and can be found in **Table S5**.

### Metabolic annotation

To estimate the metabolic potential of genera *Rhodoferax* and *Allorhodoferax*, we selected the 10 species-clusters with the largest number of genomes (i.e., species-clusters 0008, 0016, 0018, 0043, 0068, 0112, 0130, 0156, 0211 and 0339; **Table S1**).

We also selected the 3 species with the smallest average estimated genome size and at least three genomes (i.e., species-clusters 0060, 0505 and 0744) (**Table S1**). Their metabolic potential was predicted using Prodigal annotations (v2.6.3) (Hyatt et al., 2010), and the resulting protein translation files were used to predict biogeochemical and metabolic functional traits using the METABOLIC-G script from METABOLIC v4.0 (Zhou et al., 2022), using default settings. Information on the presence/absence of different metabolic genes can be found in **Table S6**.

To investigate the correlation between genome size and average completeness of biosynthetic pathways, we selected all 340 high-quality genomes (completeness > 90% and contamination < 5%) (**Table S1**). We used Anvi’o v7.1 (Eren et al., 2020) to reformat the FastA files (*anvi-script-reformat-fasta*) and create the contig databases (*anvi-gen-contigs-database*). Subsequently, we identified the KEGG pathways present in our genomes, and subset the metabolic modules for biosynthesis of amino acids and vitamins (*anvi-run-kegg-kofams* and *anvi-estimate-metabolism*) (Veseli et al., 2023). All studied modules can be found in **Table S7**.

### Metagenome collection and competitive mapping

The estimation of the relative abundance and prevalence of the 101 species-level clusters classified as genera *Rhodoferax*, *Allorhodoferax*, *Quasirhodoferax* and *Pararhodoferax* was subset from previous results (Rodríguez-Gijón et al., 2025). In summary, the representative genomes of 24,050 species-level clusters were competitively mapped against a manually curated dataset of 636 publicly available metagenomes from freshwater ecosystems (**Table S3**) using the Microbial Genomes Atlas (MiGA) v1.3.8.2 (Rodriguez-R et al., 2018) and Strobealign v0.11.0 (Sahlin, 2022), and the resulting SAM files were sorted using Samtools v1.17 (Danecek et al., 2021). To obtain the depth of each species-level cluster in our set of trimmed metagenomes, we calculated the 80% Truncated Average Depth (TAD80) using the script *BedGraph.tad.rb* v1.01 from the Enveomics collection (Rodriguez-R and Konstantinidis, 2016). To obtain the total coverage of the genomes, genome equivalents were computed using MicrobeCensus v1.1.0 (Nayfach and Pollard, 2015). Lastly, to estimate the relative abundance (%) the depth of the genomes was divided by the total depth of the microbial communities in the metagenomes. The relative abundance estimates for the four genera per metagenome can be found in **Table S4**.

### Statistical analysis

Statistical averages and the corresponding standard deviation were calculated using the functions *mean* and *sd* from the R package *stats* v4.3.2 (R Core Team, 2020). To detect statistical differences between pairs of distributions (represented as boxplots), we performed Kruskal-Wallis tests (**Figure 1C** and **4A-4C**) and Wilcoxon tests (**Figure S6**) corrected with Benjamini-Hochberg, using the function *stat_compare_means* from the R package *ggpubr* v0.6.0 (Kassambara, A., 2020). Different letters and stars in the figures indicate statistical differences of the given non-parametric test (p < 0.05). In scatterplots (**Figure S4-S6**), regression equations and the correlation coefficients to test the fit of our data were calculated using the functions *stat_regline_equation* and *stat_cor* (Pearson’s correlation coefficient) from the R package *ggpubr* v0.6.0 (Kassambara, A., 2020).

## Data availability

Information about the BioProject and/or publication of all genomes classified as *Rhodoferax*, *Allorhodoferax*, *Quasirhodoferax* and *Pararhodoferax* can be found in **Table S1**. The accession numbers and taxonomy (both NCBI and GTDB) for all 804 outgroup genomes used in the phylogenomic analysis can be found in **Table S2**. The accession numbers of all 636 metagenomes used to estimate the relative abundance of all species-clusters in (Rodríguez-Gijón et al., 2025) can be found in **Table S3**, and the corresponding estimations can be found in **Table S4**. The SeqCode registry of all newly defined genera and species can be found under the SeqCode Registry Accession: seqco.de/r:sfbkhng7.

## ACKNOWLEDGEMENTS

This work was supported by SciLifeLab. The authors acknowledge support from SNIC/Uppsala Multidisciplinary Center for Advanced Computational Science for access to the UPPMAX computational infrastructure. Computational work and data handling were enabled by resources in the projects SNIC 2022/5-392, 2023/5-126 and 2023/5-379 provided by the Swedish National Infrastructure for Computing (SNIC) at UPPMAX, partially funded by the Swedish Research Council through grant agreement no. 2018-05973. The computational results presented here have been achieved partially using the LEO HPC infrastructure of the University of Innsbruck. Additional computational resources were supported by the US National Science Foundation (NSF) through the ACCESS program with allocation MCB190042.

AR-G conceptualized and designed the research project. AR-G, LMR-R and SLG refined the project idea. AR-G compiled and curated the data. AR-G and LMR-R performed the bioinformatic analyses. AR-G and VG-M performed the data analysis. AR-G drafted the first manuscript. All authors did literature searches and edited and reviewed the manuscript.

