## Supplementary Material for "Genome size variability reflects contrasting ecological strategies in *Rhodoferax* and *Allorhodoferax*"

#### SUPPLEMENTARY FIGURES, LEGENDS AND NOTES

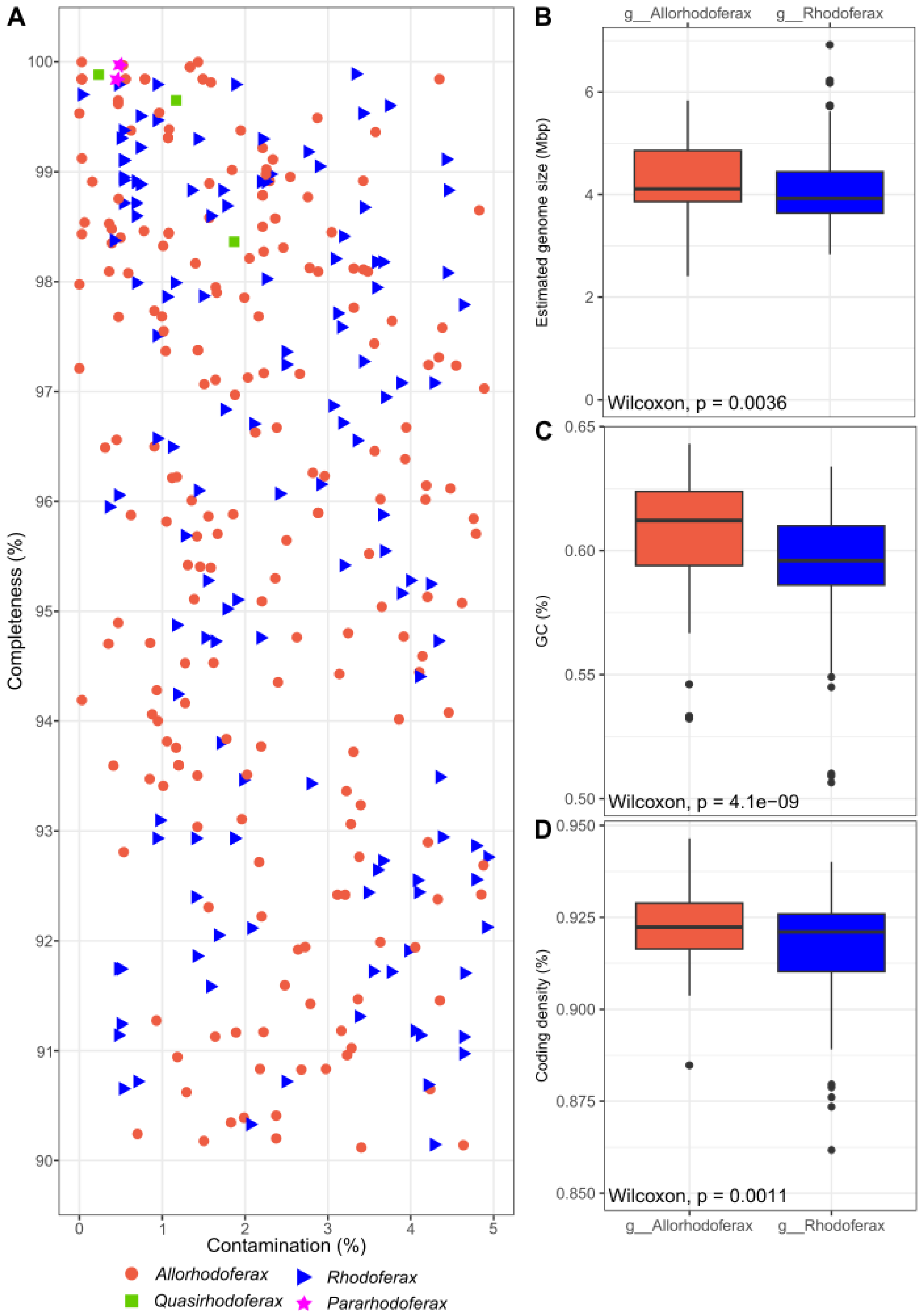


**Figure S1.** Genomic statistics all high-quality genomes included in this study. **A)** Overview of contamination and completeness per genome, indicated genera according to the legend on the bottom of the figure. We also highlight the statistical differences in estimated genome size (Mbp, B), GC content (%, C) and coding density (%, D) between *Allorhodoferax* and *Rhodoferax* genomes. Genera *Pararhodoferax* (n = 3) and *Quasirhodoferax* (n = 2) were not included given the low number of genomes in our study.

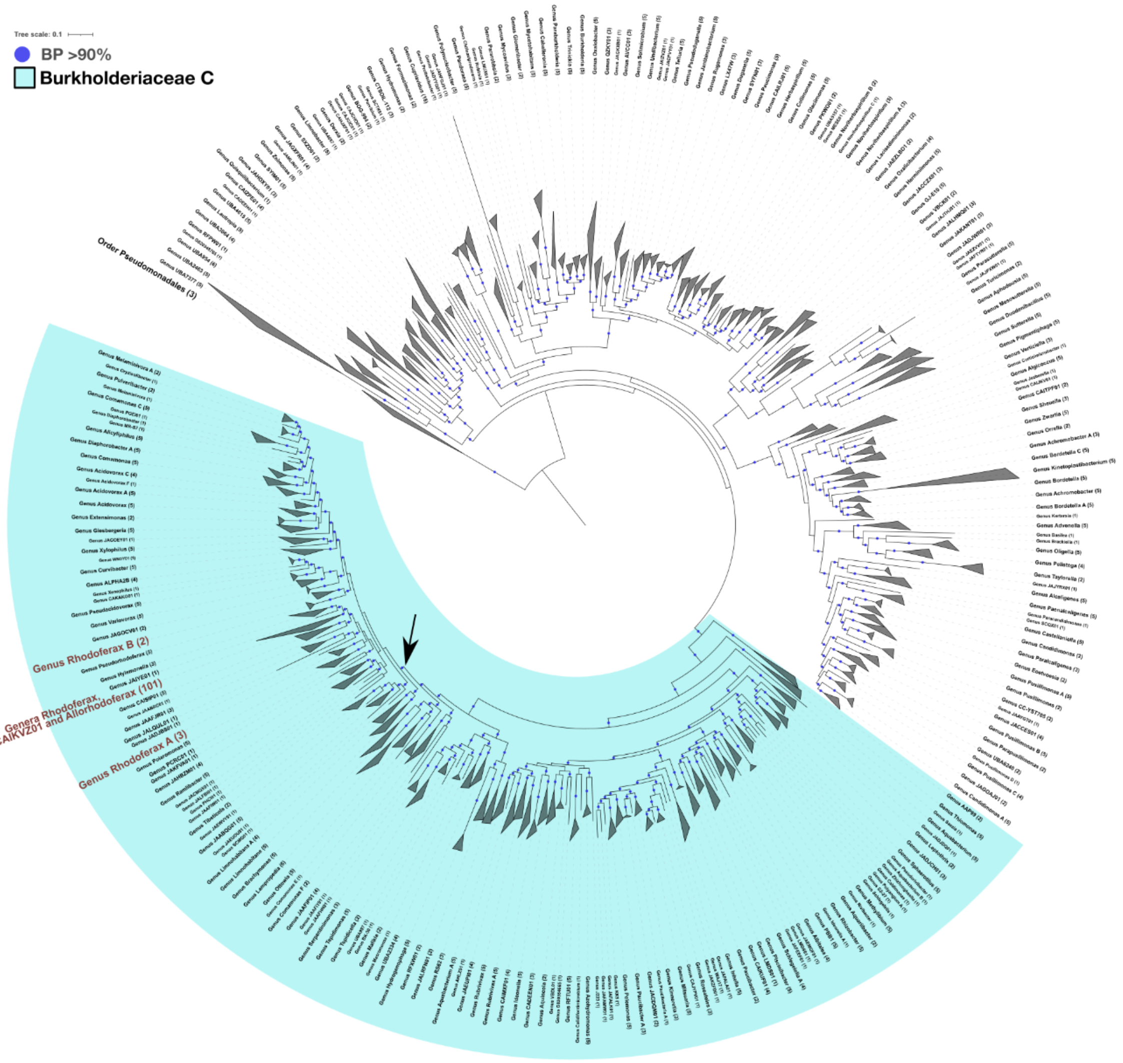


**Figure S2.** Expanded maximum-likelihood (ML) phylogeny, following a LG+F+I+I+R10 model of evolution. Phylogeny includes all 345 high-quality genomes (completeness >90% and contamination <5%) included in our study, and 801 genomes classified as *Burkholderiaceae* as outgroup. Three additional genomes from order *Pseudomonadales* were included for rooting. Accession numbers and taxonomy of root and outgroup genomes can be found in **Table S2**. Blue circles in branches indicate ≥90% bootstrap support. Arrow indicates the subset used to generate **Figure 1A**.

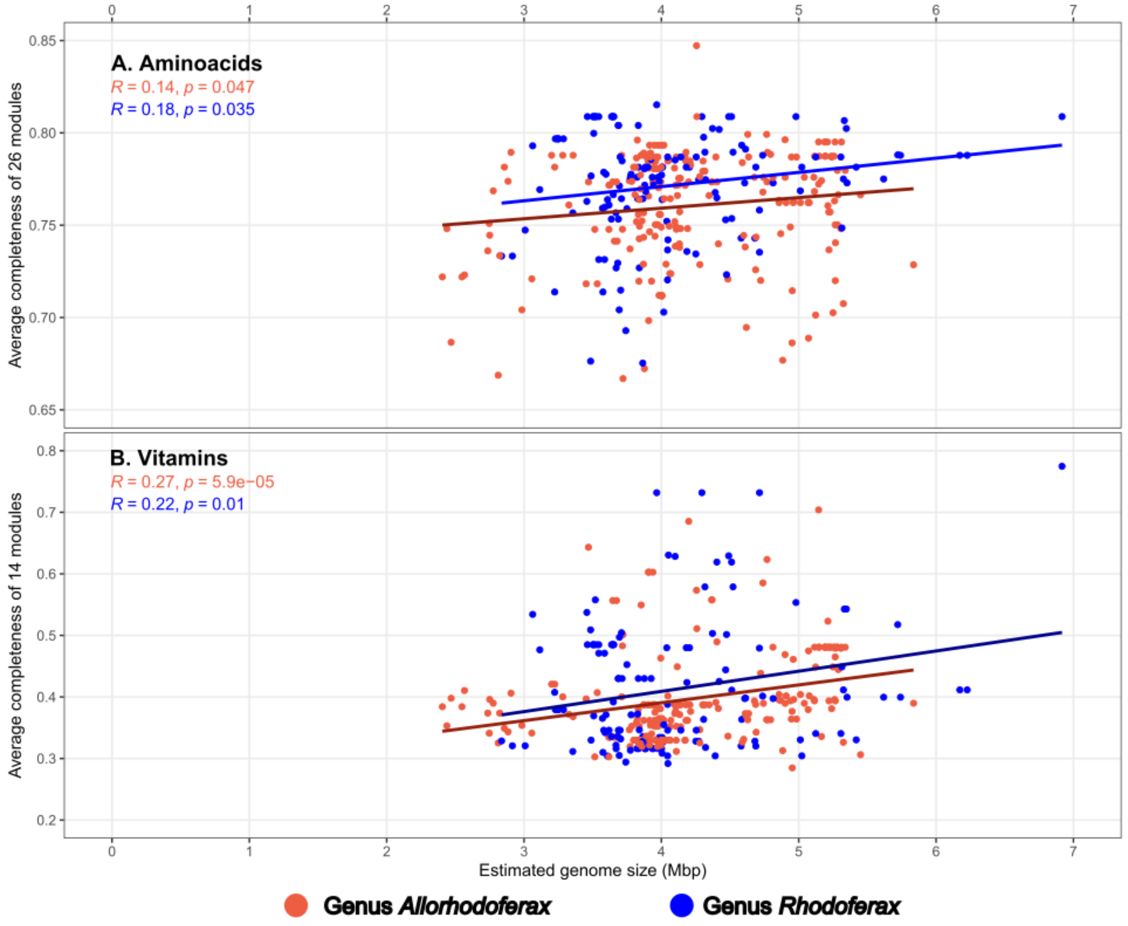


**Figure S3.** Metabolic potential for essential amino acids (**A**) and vitamins (**B**) across all 340 high-quality genomes (completeness >90% and contamination <5%) classified as genera *Rhodoferax* (n = 131) and *Allorhodoferax* (n = 209).

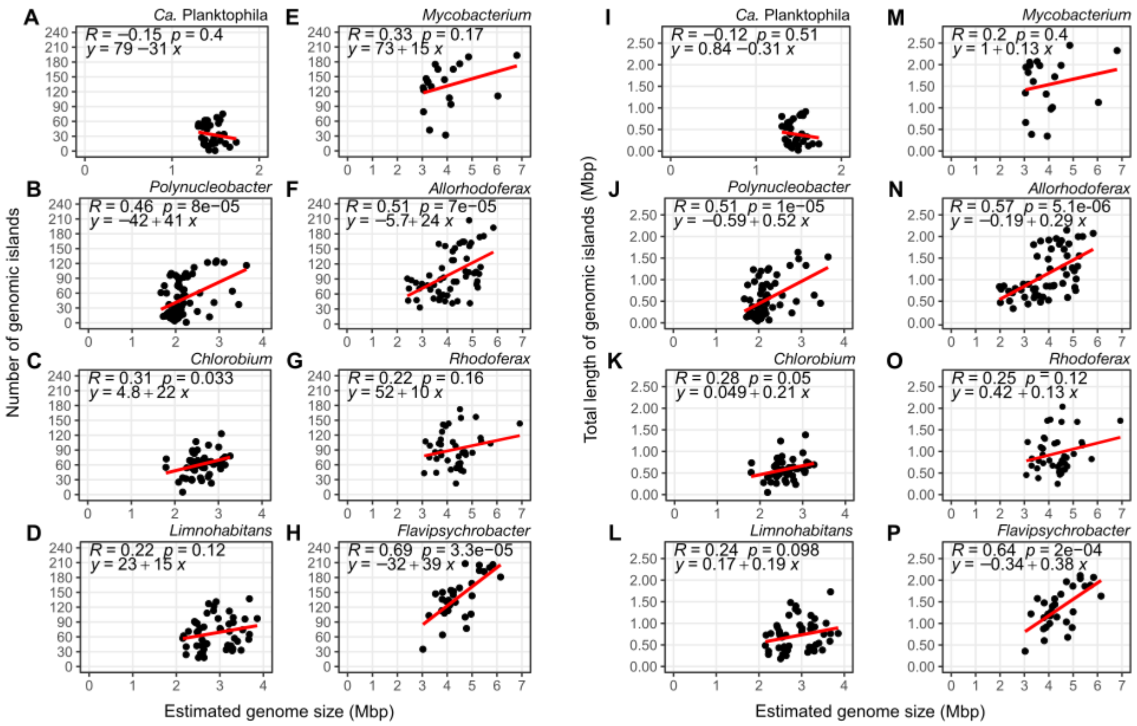

**Figure S4.** Intragenus contribution of genomic islands (GIs) to estimated genome size variability for eight freshwater bacterial genera, including genera *Allorhodoferax* and *Rhodoferax*. We overview the correlation between estimated genome size per genus (Mbp) and the number of GIs (**A**) and total length of GIs (**B**).

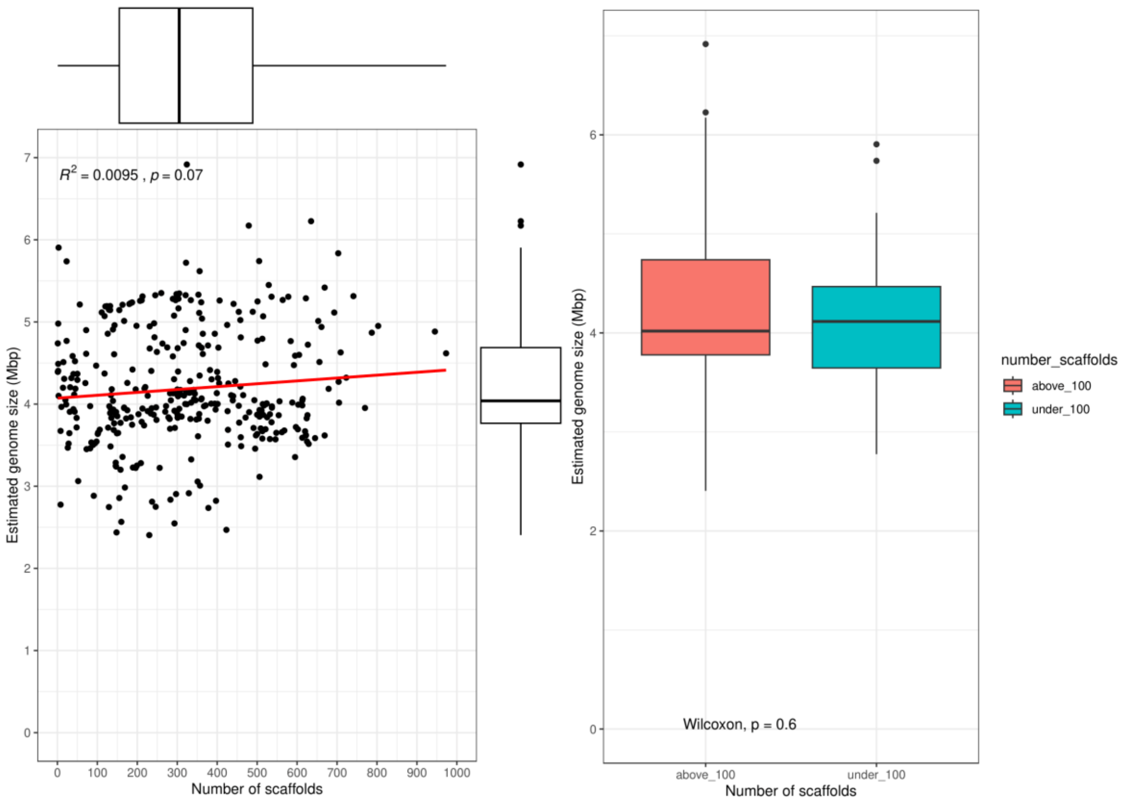


**Figure S5.** Relation between estimated genome size and the number of scaffolds across all 345 high-quality genomes (completeness >90% and contamination <5%). Both scatterplot and boxplot show non-significant correlation between the estimated genome size and the number of scaffolds.

### Register list for 17 new names including Allorhodoferax abundans sp. nov.

**Submitted by Rodríguez-Gijón, Alejandro**

#### Genus *Pararhodoferax*

**Etymology**

[Pa.ra.rho.do.fe'rax] **Gr. prep.** *para*, beside, near; **N.L. masc. n.** *Rhodoferax*, a bacterial genus; **N.L. masc. n.** *Pararhodoferax*, beside *Rhodoferax*

**Nomenclatural type**

Species *Pararhodoferax koreensis*$^{Ts}$

**Description**

The genus *Pararhodoferax* is supported by a maximum likelihood tree (bootstrap >90%) using 172 gammaproteobacterial single-copy marker genes, as well as ANI and AAI pairwise similarity analyses. This genus includes the lineage “Rhodoferax B koreense” (strain DCY-110; GCF_001955695.1, **genus type material**). The similarity of this genome in respect of *Rhodoferax fermentans* JCM 7819 (GCF_002017865.1; type strain of the genus *Rhodoferax*) is 66.08% AAI and 78.36% ANI. Genus *Pararhodoferax* also includes the genomes “Rhodoferax B 1321” (GCF_900104385.1).

Genus *Pararhodoferax* was detected in 3 of the 636 freshwater metagenomes: Långsjön lake in Sweden (SAMEA110382279), and Kalamas River in Greece (SRX1528728 and SRX1528731). Moreover, the original publications of the genomes classified as *Pararhodoferax* indicate its presence in sludge from Gangwon province in South Korea (Farh et al., 2017; Microbial Systematics and Evolutionary Microbiology 55, 767-774) and from in the rhizosphere of *Populus* sp. trees in Tennessee (US) (Nayfach et al., 2021; Nature Biotechnology 39, 499-509).

**Classification**

*Bacteria* » *Pseudomonadota* » *Betaproteobacteria* » *Burkholderiales* » *Burkholderiaceae* » *Pararhodoferax*

**Registry URL**

https://seqco.de/i:56160

### Genus *Quasirhodoferax*

**Etymology**

[Qua.si.rho.do.fe'rax] **L. conj.** *quasi*, almost, resembling but not actually being; **N.L. masc. n.** *Rhodoferax*, a bacterial genus; **N.L. masc. n.** *Quasirhodoferax*, almost *Rhodoferax*

**Nomenclatural type**

Species *Quasirhodoferax sediminis*$^{Ts}$

**Description**

The genus *Quasirhodoferax* is supported by a maximum likelihood tree (bootstrap >90%) using 172 gammaproteobacterial single-copy marker genes, as well as ANI and AAI pairwise similarity analyses. This genus includes the lineage "Rhodoferax A sediminis" (strain CHu59-6-5T; GCF_006970865.1, **genus type material**). The similarity of this genome in respect of *Rhodoferax fermentans* JCM 7819 (GCF_002017865.1; type strain of the genus *Rhodoferax*) is 66.75% AAI and 78.71% ANI. Genus *Quasirhodoferax* also includes the genomes "Rhodoferax A 1290" (GCA_004323695.1), and "Rhodoferax A 0200" (GCF_002002705.1).

Genus *Quasirhodoferax* was not detected in any of the 636 freshwater metagenomes. Yet, the original publications of the genomes here included within *Quasirhodoferax* indicate its presence in freshwater sediments from Daechung Reservoir (South Korea) (Jin et al., 2020; Microorganisms 8, 262), groundwaters from Tennessee (US) (Tian et al., 2020; Microbiome 8, 51), and acidic soils from Norway (Li et al., 2021; Nucleic Acids Research 49, D1020-D1028).

**Classification**

*Bacteria* » *Pseudomonadota* » *Betaproteobacteria* » *Burkholderiales* » *Burkholderiaceae* » *Quasirhodoferax*

**Registry URL**

https://seqco.de/i:56159

### Species *Allorhodoferax abundans*

**Etymology**

[a.bun'dans] **L. masc. part. adj.** *abundans*, abundant, referring to its high abundance

**Nomenclatural type**

Genome sc|0006138 $^{Ts}$

**Description**

Referred in our manuscript as *Allorhodoferax* 0043. *Allorhodoferax abundans* is supported by the maximum likelihood tree (bootstrap >90%; **Figure 1A**), as well as ANI and AAI pairwise similarity analyses (**Figure 1B** and **Table S1**). The proposed type genome for the species is GCA_903886275.1 (referred as Day6-2_megahit_metabat_bin-221 in **Table S1**). The similarity of this genome in respect of *Allorhodoferax saidebanchensis* ED16T (GCF_000498435.1; type strain of the genus *Allorhodoferax*) is 76.29% ANI. *Allorhodoferax abundans* was detected in 23 of the 636 freshwater metagenomes. Our results indicate its presence in different freshwater lakes, such the lakes Lomtjärnan, Erken, Långsjön, Kiruna Ki1 and Kiruna 754378–169136 (Sweden), the lakes Mekkojärvi, Alinen Mustajärvi and Ylinen Rajajärvi (Finland), the lake La Plata (Puerto Rico), the lake Toolik and the McNutt Creek (USA), and different ponds in Canada.

**Classification**

*Bacteria* » *Pseudomonadota* » *Betaproteobacteria* » *Burkholderiales* » *Comamonadaceae* » *Allorhodoferax* » *Allorhodoferax abundans*

**Registry URL**

https://seqco.de/i:56326

#### Species *Allorhodoferax loclatensis*

**Etymology**

[lo.cla.ten'sis] **N.L. masc. n.** *loclatensis*, of Loclat, referring to lake Loclat (Switzerland), location where type material was assembled

**Nomenclatural type**

Genome sc|0006480 Ts

**Description**

Referred in our manuscript as *Allorhodoferax* 0339. *Allorhodoferax loclatensis* is supported by the maximum likelihood tree (bootstrap >90%; **Figure 1A**), as well as ANI and AAI pairwise similarity analyses (**Figure 1B** and **Table S1**). The proposed type genome for the species is GCA_XXXX (referred as Loc090519-4m_bin-41 in **Table S1**. The similarity of this genome in respect of *Allorhodoferax saidebanchensis* ED16T (GCF_000498435.1; type strain of the genus *Allorhodoferax*) is 76.31% ANI. *Allorhodoferax loclatensis* was detected in 44 of the 636 freshwater metagenomes. Our results indicate its presence in different freshwater lakes, such as lake Loclat (Switzerland), lakes Erken, Lomtjärnan, Bengtsgolen, Lomtjärnan and Kiruna 754378–169136 (Sweden), lake El Tobar (Spain), lakes Mekkojärvi, Alinen Mustajärvi and Ylinen Rajajärvi (Finland), the river Elkhorn Slough (USA), and different ponds in Canada.

**Classification**

*Bacteria* » *Pseudomonadota* » *Betaproteobacteria* » *Burkholderiales* » *Comamonadaceae* » *Allorhodoferax* » *Allorhodoferax loclatensis*

**Registry URL**

https://seqco.de/i:56303

#### Species *Allorhodoferax mekkojarviensis*

**Etymology**

[me.kko.jar.vien.sis] **N.L. masc. n.** *mekkojarviensis*, of Mekkojärvi, referring to lake Mekkojärvi (Finland), the location where type material was assembled

**Nomenclatural type**

Genome sc|0006082 Ts

**Description**

Referred in our manuscript as *Allorhodoferax* 0505. *Allorhodoferax mekkojarviensis* is supported by the maximum likelihood tree (bootstrap >90%; **Figure 1A**), as well as ANI and AAI pairwise similarity analyses (**Figure 1B** and **Table S1**). The proposed type genome for the species is GCA_903928635.1 (in **Table S1** named MJ-2014_megahit_metabat_bin-0891). The similarity of this genome in respect of *Allorhodoferax saidebanchensis* ED16T (GCF_000498435.1; type strain of the genus *Allorhodoferax*) is 75.12% ANI. *Allorhodoferax mekkojarviensis* was detected in 56 of the 636 freshwater metagenomes. Our results indicate its presence in arctic ponds (northern Canada), lakes Mustajärvet, Miekojärvi, Ylinen Rajajärvi and Valkea Kotinen (Finland), lakes Lillsjon, Kiruna Ki1, Kiruna 754378–169136 (Sweden), the Rimov Reservoire (Czechia), the river Das Velhas (Brasil), the lake Monchio delle Corti (Italy) and the lake Eidfjordvatnet (Norway).

**Classification**

*Bacteria* » *Pseudomonadota* » *Betaproteobacteria* » *Burkholderiales* » *Comamonadaceae* » *Allorhodoferax* » *Allorhodoferax mekkojarviensis*

**Registry URL**

https://seqco.de/i:56276

### Species *Allorhodoferax omnivivens*

**Etymology**

[om.ni.vi.vens] **L. masc. adj.** *omnis*, everything, all; **L. pres. part.** *vivens*, living; **L. masc. part. adj.** *omnivivens*, living in any place

**Nomenclatural type**

Genome sc|0006137 [Ts]

**Description**

Referred in our manuscript as *Allorhodoferax* 0744. *Allorhodoferax ubique* is supported by the maximum likelihood tree (bootstrap >90%; **Figure 1A**), as well as ANI and AAI pairwise similarity analyses (**Figure 1B** and **Table S1**). The proposed type genome for the species is GCA_945861345.1 (**Table S1**). The similarity of this genome in respect of *Allorhodoferax saidebanchensis* ED16T (GCF_000498435.1; type strain of the genus *Allorhodoferax*) is 74.10% ANI. *Allorhodoferax ubique* was detected in 101 of the 636 freshwater metagenome. Our results indicate its presence across continents, including Asia, Europe and America. Some water bodies where A. ubique was present are the lakes Nästjärnen, Vallingasjon and Odrolstjärnen (Sweden), lakes Lovonjarvi and Haukijärvi (Finland), lake Zurich (Switzerland), the Rimov Reservoir (Czechia), lake Lanier (USA), lake Biwa (Japan) and the Narmada River (India), amongst others.

**Classification**

*Bacteria* » *Pseudomonadota* » *Betaproteobacteria* » *Burkholderiales* » *Comamonadaceae* » *Allorhodoferax* » *Allorhodoferax omnivivens*

**Registry URL**

https://seqco.de/i:56282

### Species *Allorhodoferax septentrionalis*

**Etymology**

[sep.ten.tri.o.na'lis] **L. masc. adj.** *septentrionalis*, northern

**Nomenclatural type**

Genome sc|0006481 [Ts]

**Description**

Referred in our manuscript as *Allorhodoferax* 0016. *Allorhodoferax septentrionalis* is supported by the maximum likelihood tree (bootstrap >90%; **Figure 1A**), as well as ANI and AAI pairwise similarity analyses (**Figure 1B** and **Table S1**). The proposed type genome for the species is GCA_XXX (referred as AM-lipid-02-D5_bin-13 in **Table S1**). The similarity of this genome in respect of *Allorhodoferax saidebanchensis* ED16T (GCF_000498435.1; type strain of the genus *Allorhodoferax*) is 76.40% ANI. *Allorhodoferax septentrionalis* was detected in 132 of the 636 freshwater metagenomes. Our results indicate its presence in lakes from the global north including the lakes Erken, Lomtjärnan, Malstasjön, Långsjön and Odrolstjärnen (Sweden), lakes Mekkojärvi, Alinen Mustajärvi, Ylinen Rajajärvi, Lovojärvi and Valkea Kotinen (Finland), the McNutt Creek (USA) and ponds in Canada, amongst others.

**Classification**

*Bacteria* » *Pseudomonadota* » *Betaproteobacteria* » *Burkholderiales* » *Comamonadaceae* » *Allorhodoferax* » *Allorhodoferax septentrionalis*

**Registry URL**

https://seqco.de/i:56302

#### Species *Pararhodoferax koreensis*$^{Ts}$

**Etymology**
[ko.re.en'sis] **N.L. masc. n.** *koreensis*, related to the Republic of Korea, origin of this species

**Nomenclatural type**
Genome sc|0006205 $^{Ts}$

**Classification**
*Bacteria* » *Pseudomonadota* » *Betaproteobacteria* » *Burkholderiales* » *Burkholderiaceae* » *Pararhodoferax* » *Pararhodoferax koreensis*$^{Ts}$

**Registry URL**
https://seqco.de/i:56437

#### Species *Quasirhodoferax sediminis*$^{Ts}$

**Etymology**
[se.di.mi'nis] **L. gen. n.** *sediminis*, of sediment

**Nomenclatural type**
Genome sc|0006206 $^{Ts}$

**Classification**
*Bacteria* » *Pseudomonadota* » *Betaproteobacteria* » *Burkholderiales* » *Burkholderiaceae* » *Quasirhodoferax* » *Quasirhodoferax sediminis*$^{Ts}$

**Registry URL**
https://seqco.de/i:56438

#### Species *Rhodoferax ahuizotli*

**Etymology**
[a.hu.i.zo'tli] **N.L. gen. n.** *ahuizotli*, of Ahuízotl, an aquatic creature in Mexican mythology.

**Nomenclatural type**
Genome sc|0006482 $^{Ts}$

**Description**
Referred in our manuscript as *Rhodoferax* 0008*. Rhodoferax ahuizotli* is supported by the maximum likelihood tree (bootstrap >90%; **Figure 1A**), as well as ANI and AAI pairwise similarity analyses (**Figure 1B** and **Table S1**). The proposed type genome for the species is GCA_XXX (referred as LJ-7-9m_bin-17 in **Table S1**). The similarity of this genome in respect of *Rhodoferax fermentans* JCM 7819 (GCF_002017865.1; type strain of the genus *Rhodoferax*) is 74.27% ANI. *Rhodoferax ahuizotli* was detected in 22 of the 636 freshwater metagenomes. Our results indicate its presence in lakes Lovojärvi, Mekkojärvi and Alinen Mustajärvi (Finland), lakes Odrolstjärnen and Kiruna 754378–169136 (Sweden) and the Rimov Reservoir (Czechia).

**Classification**
*Bacteria* » *Pseudomonadota* » *Betaproteobacteria* » *Burkholderiales* » *Comamonadaceae* » *Rhodoferax* » *Rhodoferax ahuizotli*

**Registry URL**
https://seqco.de/i:56427

### Species *Rhodoferax alphei*

**Etymology**
[al'phe.i] **N.L. gen. n.** *alphei*, of Alpheus, a river God in Hellenic mythology.

**Nomenclatural type**
Genome sc|0006483 Ts

**Description**
Referred in our manuscript as *Rhodoferax* 0112. *Rhodoferax alphei* is supported by the maximum likelihood tree (bootstrap >90%; **Figure 1A**), as well as ANI and AAI pairwise similarity analyses (**Figure 1B** and **Table S1**). The proposed type genome for the species is GCA_XXX (referred as Loc090402_4m_bin_2 in **Table S1**). The similarity of this genome in respect of *Rhodoferax fermentans* JCM 7819 (GCF_002017865.1; type strain of the genus *Rhodoferax*) is 74.66% ANI. *Rhodoferax alphei* was detected in 57 of the 636 freshwater metagenomes. Our results indicate its presence in the Rimov Reservoir (Czechia), the lake Biwa (Japan), the Kalamas River (Greece), the lakes Loclat and Rotsee (Switzerland), and the lakes Lumpen and Malstasjön (Sweden).

**Classification**
*Bacteria* » *Pseudomonadota* » *Betaproteobacteria* » *Burkholderiales* » *Comamonadaceae* » *Rhodoferax* » *Rhodoferax alphei*

**Registry URL**
https://seqco.de/i:56426

### Species *Rhodoferax atabeyae*

**Etymology**
[a.ta.bey'ae] **N.L. gen. n.** *atabeyae*, of Atabey, an ancestral spirit in Taíno folklore.

**Nomenclatural type**
Genome sc|0006484 Ts

**Description**
Referred in our manuscript as *Rhodoferax* 0130. *Rhodoferax atabeyae* is supported by the maximum likelihood tree (bootstrap >90%; **Figure 1A**), as well as ANI and AAI pairwise similarity analyses (**Figure 1B** and **Table S1**). The proposed type genome for the species is GCA_XXXX (referred as MJ130916A_bin-18 in **Table S1**). The similarity of this genome in respect of *Rhodoferax fermentans* JCM 7819 (GCF_002017865.1; type strain of the genus *Rhodoferax*) is 74.77% ANI. *Rhodoferax atabeyae* was detected in 113 of the 636 freshwater metagenomes. Our results indicate its presence in lakes Mekkojärvi, Keskinen Rajajärvi, Ylinen Rajajärvi, Lovojärvi and Alinen Mustajärvi (Finland), lakes Lillsjon, Odrolstjärnen, Nästjärnen, Bengtsgolen, Erken, Kiruna Ki1 and Kiruna 754378–169136 (Sweden), lake La Plata (Puerto Rico), and different ponds in Canada.

**Classification**
*Bacteria* » *Pseudomonadota* » *Betaproteobacteria* » *Burkholderiales* » *Comamonadaceae* » *Rhodoferax* » *Rhodoferax atabeyae*

**Registry URL**
https://seqco.de/i:56430

### Species *Rhodoferax memphrei*

**Etymology**

[mem'phre.i] **N.L. gen. n.** *memphrei*, of Memphré, a lake creature in Canadian folklore.

**Nomenclatural type**

Genome sc|0006201 Ts

**Description**

Referred in our manuscript as *Rhodoferax* 0068*. Rhodoferax memphrei* is supported by the maximum likelihood tree (bootstrap >90%; **Figure 1A**), as well as ANI and AAI pairwise similarity analyses (**Figure 1B** and **Table S1**). The proposed type species for the genus is GCA_903852405.1 (referred as E3_megahit_metabat_bin-010 in **Table S1**). The similarity of this genome in respect of *Rhodoferax fermentans* JCM 7819 (GCF_002017865.1; type strain of the genus *Rhodoferax*) is 74.83% ANI. *Rhodoferax memphrei* was detected in 20 of the 636 freshwater metagenomes. Our results indicate its presence lakes Mekkojärvi and Lovojärvi (Finland), lakes Lillsjon and Kiruna 754378–169136 (Sweden) and different ponds in Canada.

**Classification**

*Bacteria* » *Pseudomonadota* » *Betaproteobacteria* » *Burkholderiales* » *Comamonadaceae* » *Rhodoferax* » *Rhodoferax memphrei*

**Registry URL**

https://seqco.de/i:56429

### Species *Rhodoferax nerthi*

**Etymology**

[ner'thi] **N.L. gen. n.** *nerthi*, of Nerthus, a goddess in Germanic paganism.

**Nomenclatural type**

Genome sc|0006485 Ts

**Description**

Referred in our manuscript as *Rhodoferax* 0211*. Rhodoferax nerthi* is supported by the maximum likelihood tree (bootstrap >90%; **Figure 1A**), as well as ANI and AAI pairwise similarity analyses (**Figure 1B** and **Table S1**). The proposed type genome for the species is GCA_XXXX (referred as Loc090519-7m_bin-6 in **Table S1**). The similarity of this genome in respect of *Rhodoferax fermentans* JCM 7819 (GCF_002017865.1; type strain of the genus *Rhodoferax*) is 77.12% ANI. *Rhodoferax nerthi* was detected in 23 of the 636 freshwater metagenomes. Our results indicate its presence in lakes Rotsee and Loclat (Switzerland).

**Classification**

*Bacteria* » *Pseudomonadota* » *Betaproteobacteria* » *Burkholderiales* » *Comamonadaceae* » *Rhodoferax* » *Rhodoferax nerthi*

**Registry URL**

https://seqco.de/i:56425

### Species *Rhodoferax nixeae*

**Etymology**

[ni'xe.ae] **N.L. gen. n.** *nixeae*, of Nixe, a mythological figure in German folklore

**Nomenclatural type**

Genome sc|0006141 Ts

**Description**

Referred in our manuscript as *Rhodoferax* 0060. *Rhodoferax nixeae* is supported by the maximum likelihood tree (bootstrap >90%; **Figure 1A**), as well as ANI and AAI pairwise similarity analyses (**Figure 1B** and **Table S1**). The proposed type species for the genus is GCA_903929815.1 (referred as Loclat_megahit_metabat_bin-01332 in **Table S1**). The similarity of this genome in respect of *Rhodoferax fermentans* JCM 7819 (GCF_002017865.1; type strain of the genus *Rhodoferax*) is 70.78% ANI. *Rhodoferax nixeae* was detected in 20 of the 636 freshwater metagenomes. Our results indicate its presence in only three freshwater lakes: the lakes Loclat and Rotsee (Switzerland) and the lake Lomtjärnan (Sweden).

**Classification**

*Bacteria* » *Pseudomonadota* » *Betaproteobacteria* » *Burkholderiales* » *Comamonadaceae* » *Rhodoferax* » *Rhodoferax nixeae*

**Registry URL**

https://seqco.de/i:56329

### Species *Rhodoferax sjoraaetae*

**Etymology**

[sjo.raa'et.ae] **N.L. gen. n.** *sjoraaetae*, of Sjörået, a mythological figure in Norse folklore

**Nomenclatural type**

Genome sc|0006200 Ts

**Description**

Referred in our manuscript as *Rhodoferax* 0156. *Rhodoferax sjoraaetae* is supported by the maximum likelihood tree (bootstrap >90%; **Figure 1A**), as well as ANI and AAI pairwise similarity analyses (**Figure 1B** and **Table S1**). The proposed type species for the genus is GCA_903823025.1 (referred as Day6-1_megahit_metabat_bin-090 in **Table S1**). The similarity of this genome in respect of *Rhodoferax fermentans* JCM 7819 (GCF_002017865.1; type strain of the genus *Rhodoferax*) is 73.24% ANI. *Rhodoferax sjoraaetae* was detected in 12 of the 636 freshwater metagenomes. Our results indicate its presence in only one freshwater lake, the lake Lomtjärnan (Sweden).

**Classification**

*Bacteria* » *Pseudomonadota* » *Betaproteobacteria* » *Burkholderiales* » *Comamonadaceae* » *Rhodoferax* » *Rhodoferax sjoraaetae*

**Registry URL**

https://seqco.de/i:56424

### Species *Rhodoferax urdarbrunnri*

**Etymology**

[ur.dar.brun'nri] **N.L. gen. n.** *urdarbrunnri*, of Urðarbrunnr, a sacred well in Norse mythology.

**Nomenclatural type**

Genome sc|0006486 Ts

**Description**

Referred in our manuscript as *Rhodoferax* 0018. *Rhodoferax urdarbrunnri* is supported by the maximum likelihood tree (bootstrap >90%; **Figure 1A**), as well as ANI and AAI pairwise similarity analyses (**Figure 1B** and **Table S1**). The proposed type genome for the species is GCA_XXXX (referred as MJ130916A_bin-5 in **Table S1**). The similarity of this genome in respect of *Rhodoferax fermentans* JCM 7819 (GCF_002017865.1; type strain of the genus *Rhodoferax*) is 74.84% ANI. *Rhodoferax urdarbrunnri* was detected in 80 of the 636 freshwater metagenomes. Our results indicate its presence in lakes Alinen Mustajärvi, Mekkojärvi, Keskinen Rajajärvi, Valkea Kotinen and Ylinen Rajajärvi (Finland), and lake Erken (Sweden).

**Classification**

*Bacteria* » *Pseudomonadota* » *Betaproteobacteria* » *Burkholderiales* » *Comamonadaceae* » *Rhodoferax* » *Rhodoferax urdarbrunnri*

**Registry URL**

https://seqco.de/i:56428
